# Origins of Scaling Laws in Microbial Dynamics

**DOI:** 10.1101/2021.05.24.445465

**Authors:** Xu-Wen Wang, Yang-Yu Liu

**Affiliations:** Channing Division of Network Medicine, Department of Medicine, Brigham and Women’s Hospital and Harvard Medical School, Boston, MA, 02115, USA

## Abstract

Analysis of high-resolution time series data from the human and mouse gut microbiomes revealed that the gut microbial dynamics can be characterized by several robust and simple scaling laws. It is still unknown if those scaling laws are universal across different body sites, host species, or even free-living microbial communities. Moreover, the underlying mechanisms responsible for those scaling laws remain poorly understood. Here, we demonstrate that those scaling laws are not unique to gut microbiome, but universal across different habitats, from human skin and oral microbiome to marine plankton bacteria and eukarya communities. Since completely shuffled time series yield very similar scaling laws, we conjecture that the universal scaling laws in various microbiomes are largely driven by temporal stochasticity of the host or environmental factors. We leverage a simple population dynamics model with both deterministic inter-species interactions and stochastic noise to confirm our conjecture. In particular, we find that those scaling laws are jointly determined by inter-species interactions and linear multiplicative noises. The presented results deepen our understanding of the nature of scaling laws in microbial dynamics.

## I. INTRODUCTION

Microorganisms grow and thrive in all habitats throughout the biosphere [1–7]. Those microorganisms play a critical role in maintaining the well-being of their hosts [8–12] or the integrity of their environment [13–15]. Microbiome dysbiosis can markedly affect the host’s health status [16–18] and is associated with many diseases [19–22]. Numerous studies have demonstrated that microbiomes are considerably dynamic and can be regulated by many host and environmental factors, i.e., diet, medication and host lifestyle [23–27]. Interestingly, it has been found through comprehensive analyses of high-resolution time series data that, despite inherent complexity, the dynamics of the human and mouse gut microbiomes display several simple and robust scaling laws [28]. This finding raises two fundamental questions. First of all, are those scaling laws unique for human and mouse gut microbiomes or universal across different body sites and host species, or even free-living microbiomes? Second, what are the underlying mechanisms responsible for those scaling laws? Recently, we found that those scaling laws can still be observed from the shuffled time series of the human and mouse gut microbiomes, where the temporal structure in the original time series has been largely destroyed [29]. This finding prompts us to hypothesize that temporal fluctuations might be the key to explain those scaling laws. But this hypothesis has not been quantitatively validated yet.

To address those questions, we first analyzed high-resolution time series microbiome data from different habitats, finding that those scaling laws previously observed in the human and mouse gut microbiomes actually are universal across different habitats, regardless of being host-associated or free-living microbiome. Then, we leveraged a population dynamic model with both deterministic inter-species interactions and stochastic noise, finding that the emergence of those scaling laws is jointly determined by inter-species interactions and linear multiplicative noises.

## II. RESULT

### A. High-resolution microbiome time series data analysis

Let us consider a time series of microbial compositions *X*_*k*_(*t*) of a particular habitat. Here, *X*_*k*_(*t*) represents the relative abundance of taxon-*k, k* = 1, …, *N*, and *t* = 1, …, *T*. Several scaling laws have been proposed to describe the dynamics of human and mouse gut microbiomes, based on longitudinal 16S rRNA gene sequencing data analysis [28] (see Table.I): (1) Distribution of short-term abundance change; (2) Variability of short-term abundance change; (3) Long-term drift; (4) Residence (return) time distribution; and (5) Taylor’s law that relates the variances of species’ abundances to their means.

Here we introduce a completely new scaling law: the degree distribution of the visibility raph associated with the time series of microbiome data follows an exponential distribution: *P*(*k*) ∝ exp(−*αk*), or log *P*(*k*) ∝ −*αk*, where *k* is the degree of a taxon in the visibility graph. Transformed from time series, visibility graphs allow us to study dynamical systems through the characterization of their associated networks [30]. For example, a periodic time series can be mapped into a regular graph, a random time series can be mapped into an Erdős–Rényi random graph with a Poisson degree distribution, and a fractal time series can be mapped into a scale-free graph with a power-law degree distribution.

To check the universality of those scaling laws, we analyzed high-resolution time series data of various microbiomes, from human gut [23], skin [25], oral [25], to mouse gut microbiome [31], and marine plankton bacteria and eukarya communities [32] (see Fig.1 for the stream plots of the various time series and SI Table S1 for details of those datasets). As expected, the compositions of those microbiomes are highly dynamic over time. Then, we confirmed that the five previously reported scaling laws (Table I: laws (1)-(5)) in the human and mouse gut microbiomes can also be observed in human skin and oral microbiome, as well as the marine bacteria and eukarya communities (Fig.2, columns 1-5, red solid dots). The same is true for the new scaling law on the visibility graph degree distribution (Fig.2, column-6, red solid dots). Moreover, the exponents of most scaling laws are quite close to what have been discovered in the human and mouse gut microbiomes (see SI Table S2 for exponent values obtained from the time series of various microbiomes).

**FIG. 1:**
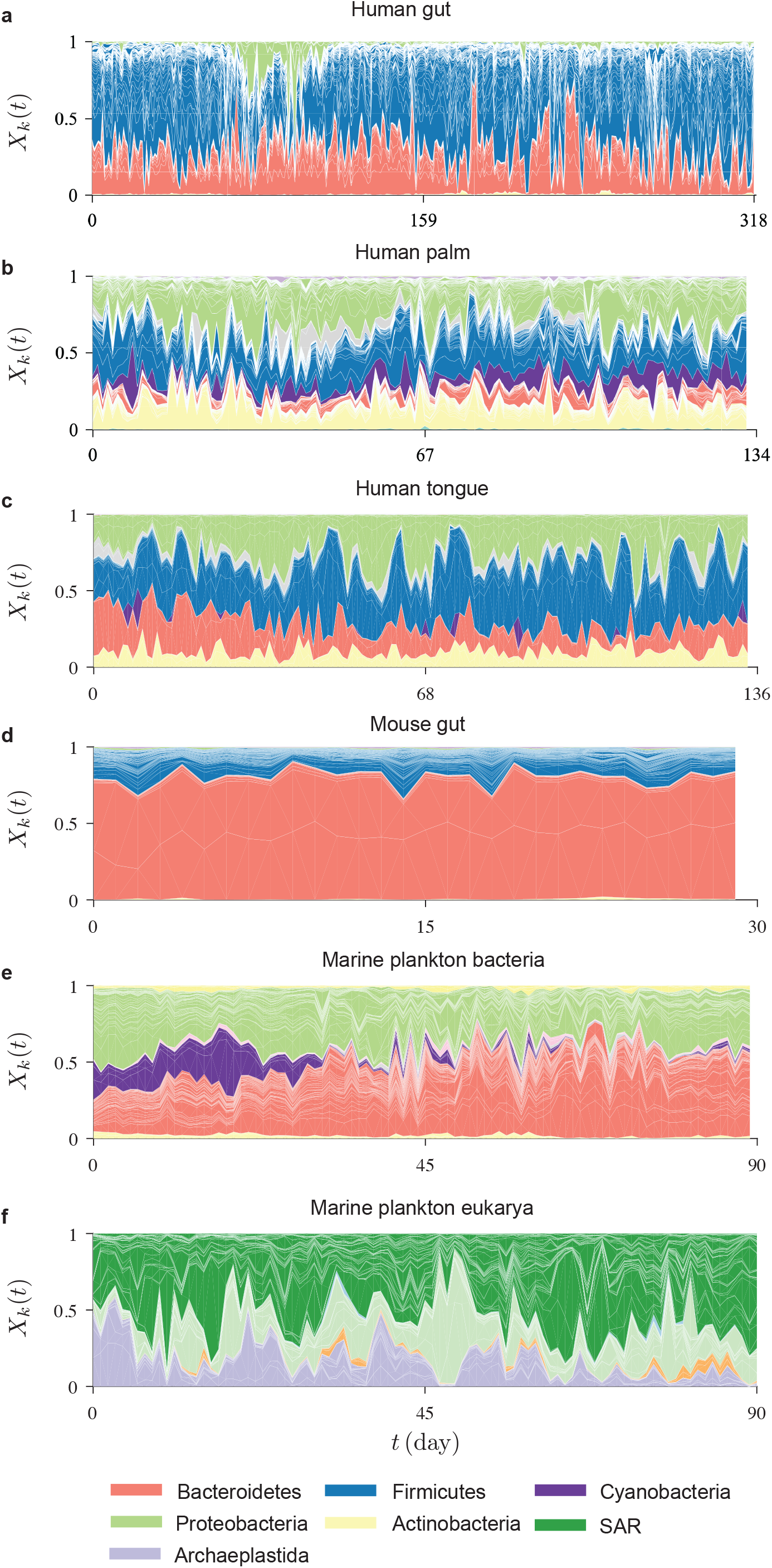
Time series of various microbiomes. (a) Human gut [23]; (b) Human palm [25]; (c) Human oral [25]; (d) Mouse gut [31]; (e) Marine plankton bacteria community and (f) Marine plankton eukarya community [32]. Stream plots showing the relative abundances of different taxa over time. Each stream represents a particular taxon and streams are grouped by phylum. Stream widths reflect the relative abundances of those taxa. For human gut, mouse gut, marine plankton bacterial and enkarya communities, the taxonomic profiles were quantified at the OTU level. For human palm and oral microbiomes, the taxonomic profiles were quantified at the genus level.

**TABLE I:**
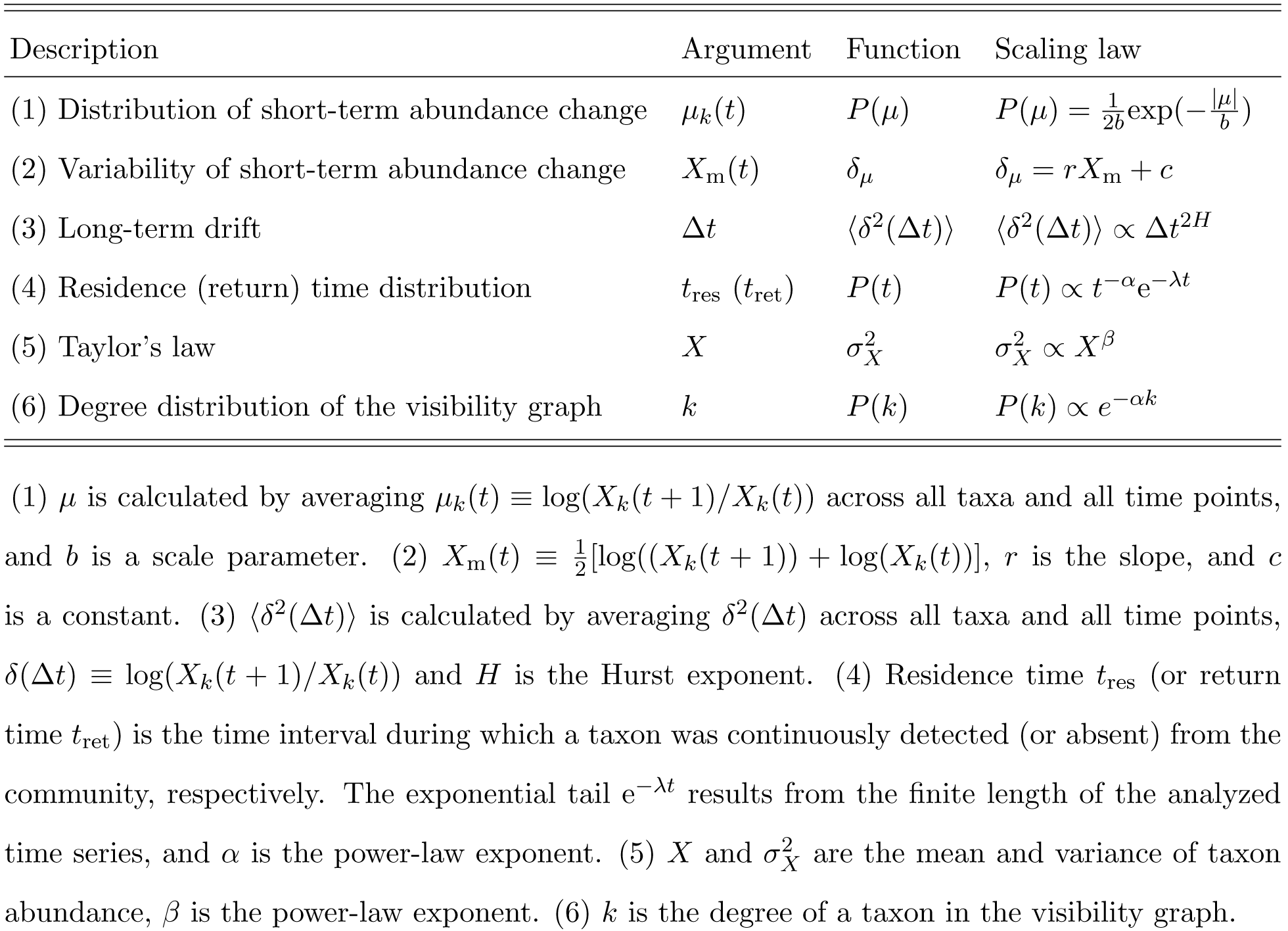
Scaling laws in microbial dynamics

To understand the nature of those scaling laws, we introduced a null model by randomly shuffling the time series to destroy the temporal structure in the original time series [29]. We found that most of the scaling laws can still be observed up to the change of the exponent values (Fig.2, blue hollow dots). For certain scaling laws (e.g., the power-law distributions of *t*_res_ and *t*_ret_, and the exponential degree distribution of the visibility graph), the shuffling will even keep the exponents almost unchanged (Fig.2, column-4, column-6). As for Taylor’s law, it will not be affected by the shuffling at all (Fig.2, column-5). In the original time-series data, the compositions of any two closely collected samples tend to be more similar to each other than samples collected with long intervals. The shuffling will significantly eliminate this impact, rendering the exponents in the scaling law of the long-term drift (Fig.2, column-3) much lower in the null model than in the original time series (see SI Table S3 for exponent values obtained from shuffled time series). Since completely shuffled time series yield similar scaling laws (up to the change of some exponent values), we conjecture that the universal scaling laws in various microbiomes are largely driven by temporal stochasticity of the host or environmental factors.

**FIG. 2:**
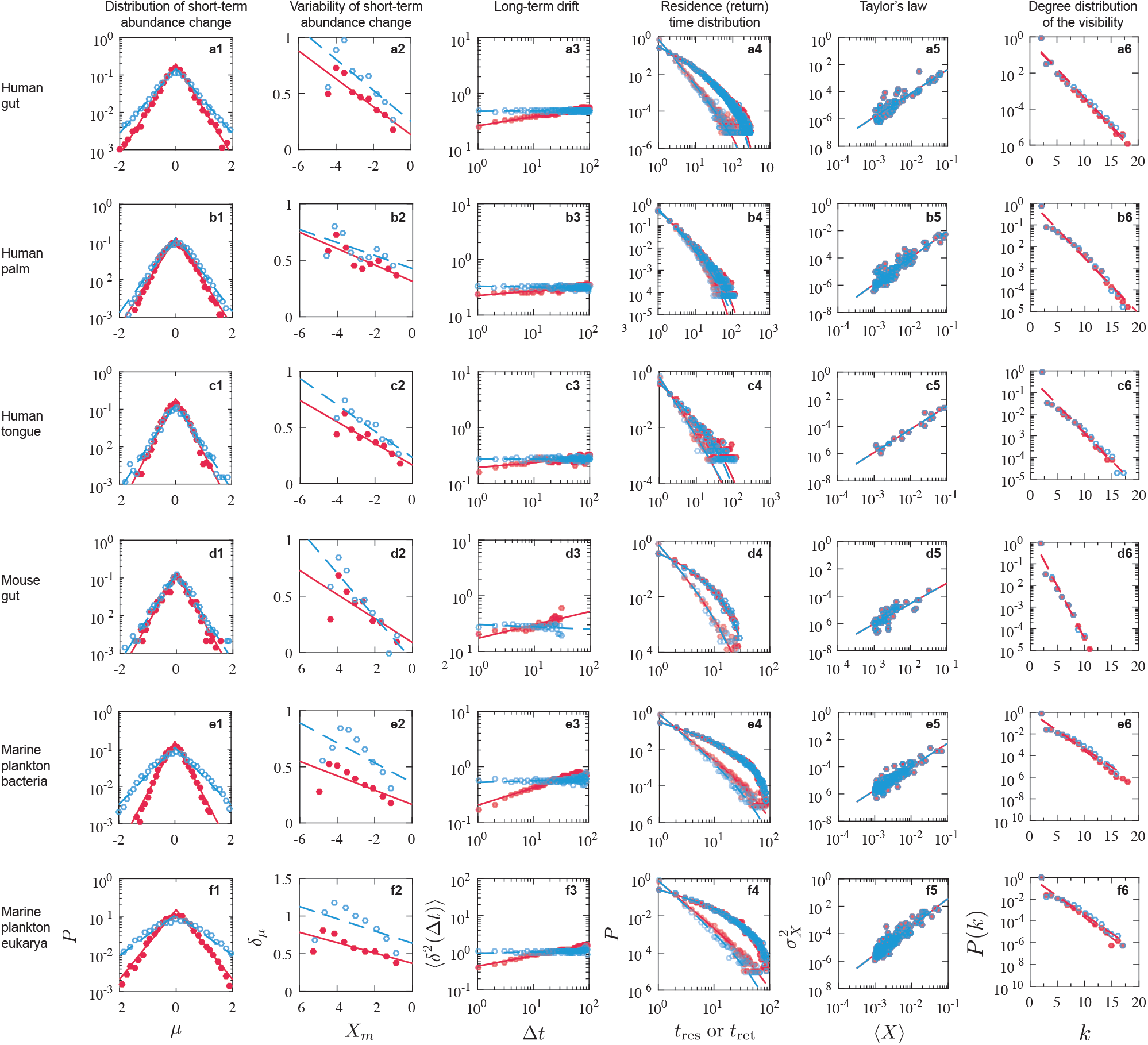
Scaling laws observed from the time series data of various microbiomes. Throughout this figure, solid (or hollow) dots represent results obtained from the original (or shuffled) time series, respectively. Lines represent maximum likelihood estimation (MLE) fits to the data. To control for known technical factors such as sample preparation and sequencing noise, here we adopted exactly the same taxa inclusion criteria as used by Refs [28, 33]. Rows: (a) Human gut; (b) Human palm; (c) Human tongue; (d) Mouse gut; (e) Marine plankton bacterial community and (f) Marine plankton eukarya community. Column: (1) Short-term (daily) abundance change *μ* follows a Laplace distribution: 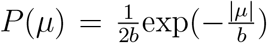. (2) The variability (i.e., standard deviation) of daily abundance change *δ* _*μ*_ decreases approximately linearly with increasing the mean of successive log-abundance *X*_m_(*t*). (3) The long-term drift of gut microbiota abundance can be approximated by the equation of anomalous: ⟨ *δ* ^2^(Δ *t*) ∝ Δ *t*^2*H*^, where *H* is the so-called Hurst exponent. The maximum lag is set to be (4) The distributions of residence (*t*_res_) and return times (*t*_ret_) follow power laws with exponential tails: *P*(*t*) ∝ *t*^*-*α^ e^*-*λ *t*^. (5) The temporal variability patterns of gut microbiota closely follow Taylor’s law: 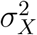, where *X* and 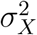 are the mean and variance of species abundance. (6) Degree distribution of the visibility graph follows an exponential distribution: *P*(*k*) ∝ exp(−*αk*).

### B. Simulations based on a simple population dynamic model

To reveal the origins of those scaling laws and check if they are largely driven by temporal stochasticity of the host or environmental factors, we added a stochastic term to the classical Generalized Lotka-Volterra (GLV) population dynamic model to incorporate external fluctuations, yielding a set of stochastic differential equations [33]:

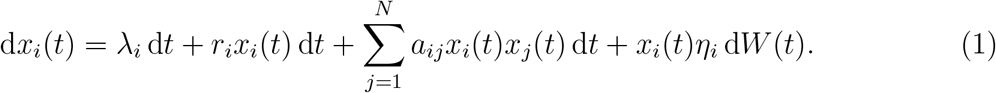

Here, *x*_*i*_, *λ* _*i*_ and *r*_*i*_ represent the abundance, immigration rate and the intrinsic growth rate of species-*i*, respectively. *N* is the total number of species. The inter-species interaction strengths are encoded in the matrix *A* = (*a*_*ij*_) ∈ ℝ ^*N*× *N*^, where *a*_*ij*_ (*i* ≠ *j*) is the per capita effect of species-*j* on the per capita growth rate of species-*i*, and *a*_*ii*_ represents the intra-species interactions. 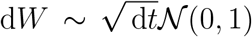 is an infinitesimal element of Brownian motion defined by a variance of d*t. η* is the noise strength. Here we consider that the stochastic term represents fluctuations of host or environmental factors, which translates into fluctuations of the intrinsic growth rate *r*_*i*_. Therefore, this term is proportional to *x*_*i*_. In other words, the stochastic term represents linear multiplicative noise.

We drew the growth rate *r*_*i*_ from a normal distribution 𝒩 (*m*, 1) and the immigration rate was set to λ = 0 (see SI Fig.S1 for the scaling laws generated with λ > 0). The diagonal elements of the interaction matrix *A* was set to be −1, while the off-diagonal elements *a*_*ij*_’s were drawn from a normal distribution 𝒩 (0, *σ*^2^) with probability *C* and set to be zero with probability (1 − *C*). Hence, *C* represents the connectivity of the underlying ecological network, and *σ*; represents the characteristic inter-species interaction strength.

In all our simulations, we let the system evolve from a random initial state (with species abundances drawn from a uniform distribution *𝒰* (0,*N*)) into the basin of a steady-state attractor. Here we assume that the temporal variations of species abundances are just reflecting fluctuations around a particular fixed point (steady state) of the dynamical system. This assumption is consistent with previous finding that the human gut microbiome can be considered as a dynamically stable ecosystem, continually buffeted by internal and external forces and recovering back toward a conserved steady-state [23]. We assume this is a general picture of various microbiomes, regardless of being host-associated or free-living.

In our modeling framework, there are four key parameters: (1) the network connectivity *C*; (2) the characteristic inter-species interaction strength *σ*; (3) the noise level *η*; and (4) the mean intrinsic growth rate *m*.

We first considered the high network connectivity and strong characteristic interaction strength: *C* = 1, *σ* = 0.1. With this fixed pair of (*C, σ*), we checked the impacts of *η* and *m* on the various scaling laws (see Fig.S2 for the stream plots of the corresponding time series). We found that for lower *m*, the short-term abundance changes tend to follow a Gaussian rather than Laplacian distribution (Fig.3a1,b1). Also, both the distribution and variability of short-term abundance changes (*P*(*μ*) and *δ*_*μ*_) after time series shuffling look quite different from those obtained from the original time series (Fig.3a1,b1; a2,b2). For lower *η* (Fig.3a1,c1), *P*(*μ*) at *μ* = 0 is much higher than the case of higher *η* (Fig.3b1,d1). This can be explained by the fact that in the presence of weak noises (i.e., low *η*), species abundance changes tend to be more deterministic than the case of strong noises (high *η*). Note that other scaling laws are not largely affected by the noise level *η*. Interestingly, with higher *m* and *η*, the stochastic GLV model can generate time series that display all the six scaling laws (Fig.3d) as observed in the real microbiome time series (Fig.2).

**FIG. 3:**
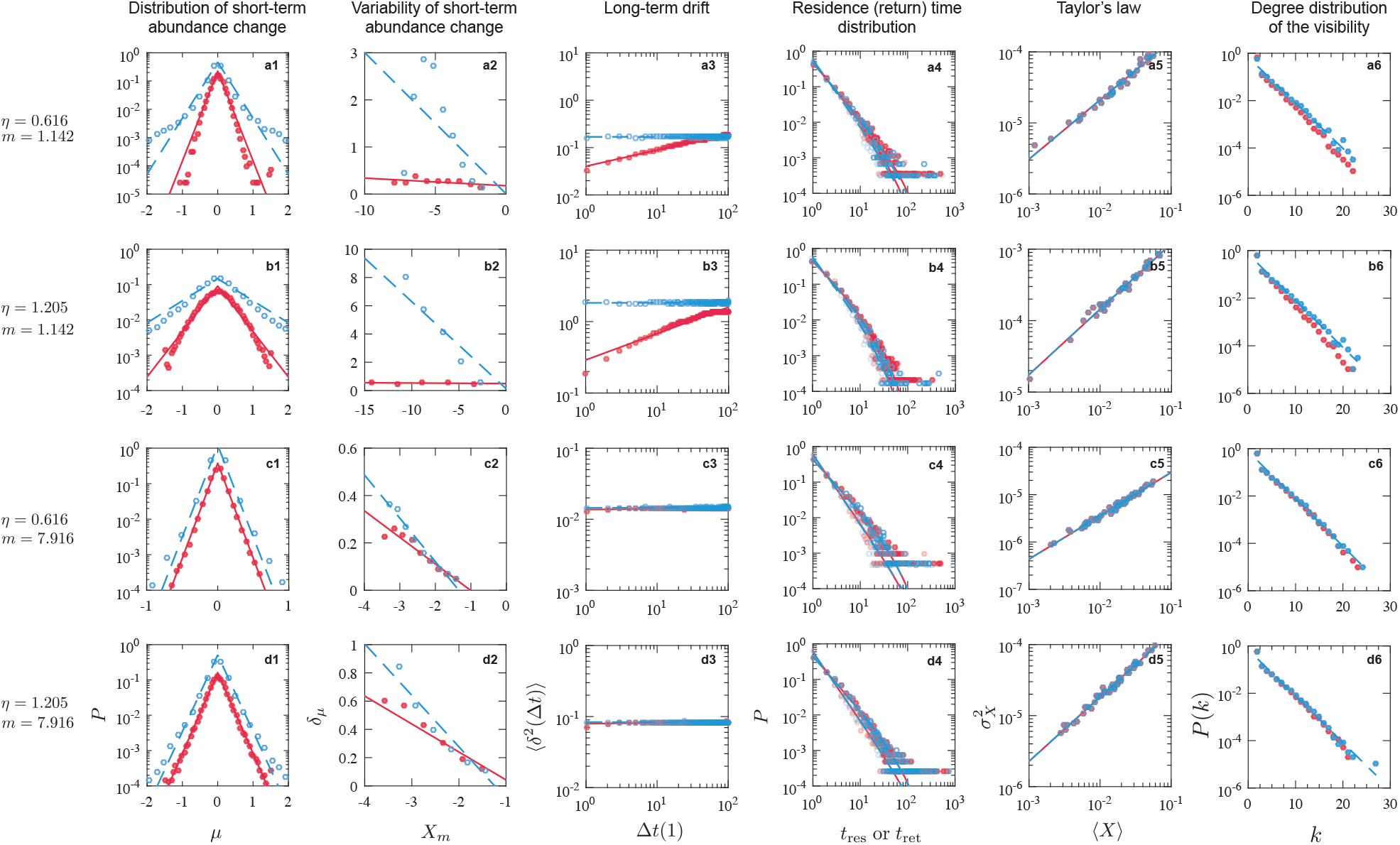
Scaling laws observed from simulated time series generated by the stochastic GLV model with various levels of noise magnitude (*η*) and mean intrinsic growth rate (*m*). Lines represent maximum likelihood estimation (MLE) fits to the data. Rows: (1) *η* = 0.616, *m* = 1.142; (2) *η* = 1.205, *m* = 1.142; (3) *η* = 0.616, *m* = 7.916 and (4) *η* = 1.205, *m* = 7.916. The columns represent those scaling laws as shown in Fig.2. *C* = 1, *σ* = 0.1 and *N* = 100.

Among all the six scaling laws, *P*(*μ*) is the one that is most sensitive to model parameters. Therefore, we next systematically checked the impacts of all the four parameters (*C, σ, η*, and *m*) on *P*(*μ*). In particular, we tried nine pairs of (*C, σ*), covering low, intermediate and high levels of network connectivity and characteristic interaction strength. For each pair of (*C, σ*), we systematically tune *η* and *m* values. For each parameter combination, we calculated 20 time series from independent stochastic GLV model instances. We then calculated *P*(*μ*) from the time series and fitted the data using both Laplacian and Gaussian distributions. We quantified the goodness of fit by the Akaike Information Criterion (AIC), calculated based on the maximum likelihood estimate (MLE). As shown in each panel (corresponding to a particular (*C, σ*) pair) of Fig.4, the color represents the probability that the AIC of the fitting with Laplacian distribution is lower than that using Gaussian distribution over the 20 independent time series data. The higher this probability the better the Laplacian distribution fits the data.

**FIG. 4:**
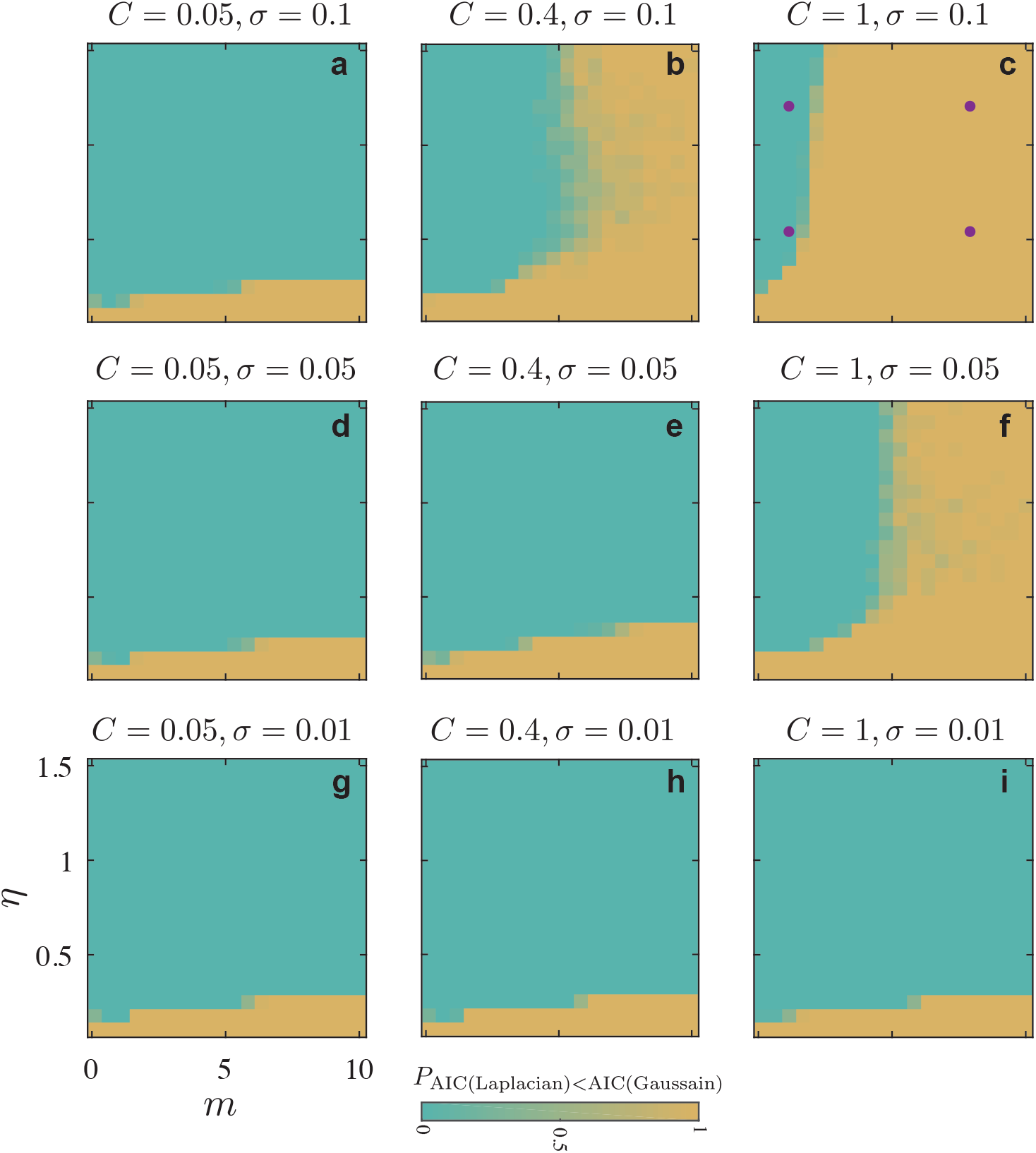
The distribution of short-term abundance change, *P*(*μ*), depends on various of parameters: *η*: the noise level, *m*: the mean of intrinsic growth rate, *σ*: the characteristic interaction strength, *C*: the connectivity of the ecological network. For each parameter combination, we generated 20 independent time series data, then fit the data using both Laplacian distribution and Gaussian distribution. The color represents the probability that the Akaike Information Criterion (AIC) calculated based on the maximum likelihood estimate (MLE) fits to the data using the Laplacian distribution is lower than that using Gaussian distribution over 20 independent time series data. Note that the higher this probability the better the Laplacian distribution fits the data. The time step is d*t* = 0.01, total time *T* = 1000, and the number of species is *N* = 100. The purple dots shown in panel-c correspond to the four (*η, m*) pairs (rows) demonstrated in Fig.3.

In a sense, each panel of Fig.4 can be regarded as a (*η, m*)-phase diagram, where *P*(*μ*) tends to behave like either a Laplacian (in the yellow region) or a Gaussian distribution (in the cyan region). We found that the shape of this (*η, m*)-phase diagram depends on the values of the (*C, σ*). When either *σ* or *C* is low, the (*η, m*)-phase diagram is dominated by the “Gaussian phase”. In this case, only when the noise level if very low, *P*(*μ*) will behave like a Laplacian distribution. Interestingly, when both *C* and *σ* are high, the (*η, m*)-phase diagram will be dominated by the “Laplacian phase”. In this case, only for very low mean intrinsic growth rate *m, P*(*μ*) will behave like a Gaussian distribution.

Note that the emergence of a Gaussian distribution of *P*(*μ*) for certain parameter settings is not a big surprise. After all, the growth of microorganisms is affected by random multiplicative processes [34, 35]. If we generalize the definition of short-term abundance change as: *μ*_*k*_(*t*)= log(*X*_*k*_(*t* + *τ*)*/X*_*k*_(*t*)), where *τ* is the length of the “short term”(and when *τ* =1 this reduces to the original definition of *μ*_*k*_(*t*)), the (*η, m*)-phase diagram will be dominated by the “Laplacian phase” with increasing *τ* (see SI Fig.S3).

It has been reported before that inter-species interactions can explain Taylor’s law for ecological time series [36]. As the variance of species abundance is largely determined by inter-species interactions, we confirmed that Taylor’s law can be reproduced regardless of the detailed values of *η* and *m* for any *C* > 0 and *σ* > 0 (see Fig.3, column-5, for results with *C* =1 and *σ* = 0.1). But in the absence of inter-species interactions (i.e., *C* = 0, *σ* = 0), we cannot reproduce Taylor’s law as observed from real data. In particular, the difference of different species’ mean abundance is very small, yielding a very concentrated scatter plot of 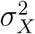 vs. ⟨*X*⟩ (see SI Fig.S4, column-5). Note that in this case, the Laplacian distribution of short-term abundance change *P*(*μ*) cannot be reproduced either. Also, the distributions of residence and return times, *P*(*t*_res_) and *P*(*t*_ret_), look quite different from what we observed from real data.

We also found that other types of noise terms, e.g., square-root multiplicative noise 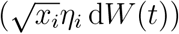 cannot reproduce the shape of Taylor’s law as observed in real data. For linear multiplicative noise, the variations of dominating species can be much higher than that of low-abundance species. By contrast, for square-root multiplicative noise, the variation difference between dominating and low-abundance species will be much smaller, yielding a very concentrated scatter plot of 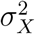 vs. ⟨*X*⟩ (see SI Fig.S5, column-5).

## III. DISCUSSION

The presented results revealed the origins of universal scaling laws in the dynamics of various microbiomes. Those scaling laws reflect the species abundance fluctuations around a stable equilibrium of the ecological system, and the fluctuations are largely driven by temporal stochasticity of the host or environmental factors. Furthermore, he presence of those scaling laws is jointly determined by inter-species interactions and linear multiplicative noise. The presented results help us better understand the nature of those universal scaling laws in the dynamics of various microbiomes.

Many of the scaling laws described here for microbial communities have also been observed previously in various macroecological systems, despite the difference of more than six orders of magnitude in the relevant spatial and interaction scales [28]. We anticipate that our simple mechanisms based on inter-species interactions and linear multiplicative noise might be universal to explain those scaling laws in both macroscopic and microbial communities.

## Acknowledgment

Research reported in this publication was supported by grants R01AI141529, R01HD093761, R01AG067744, UH3OD023268, U19AI095219, and U01HL089856 from National Institutes of Health.

## Author Contributions

Y.-Y.L. conceived and designed the project. X.-W.W. performed all the numerical calculations and data analysis. Both authors analyzed the results and wrote the manuscript.

## Author Information

The authors declare no competing financial interests. Correspondence and requests for materials should be addressed to Y.-Y.L.

## Supplementary Information

### I. MICROBIOME DATASETS

We analyzed high-resolution time series data of different microbial communities reported in Refs. [1, 2]. To control for known technical factors such as sample preparation and sequencing noise, here we adopted exactly the same taxon inclusion criteria as reported in Ref. [1, 2]. Note that the time series data of stool microbiome samples from four human subjects and six mice are available in Ref. [1]. To reduce the redundancy, we chose Human A and the first mouse with low-fat, plant polysaccharid diet in Ref. [1]. The detailed description of each dataset can be found in Table.S1.

### II. VISIBILITY GRAPH

Visibility graph allows us to transform a time series into a network. Visibility means that two data points in the time series can “see” each other without any obstacle. For any two points (*t*_*a*_, *y*_*a*_) and (*t*_*b*_, *y*_*b*_), there is a link between two nodes (or they are mutually visible), if and only if for any *t*_*c*_ (*t*_*a*_ < *t*_*c*_ < *t*_*b*_) [7, 8]:

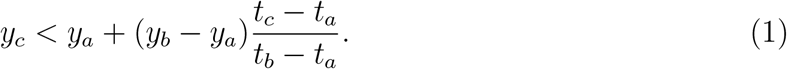

We used the Matlab implementation of the fast natural visibility graph algorithm to get the visibility graphs of the various microbiome time series [9, 10].

**TABLE S1.** Description of various microbiome time series analyzed in this study.

| Dataset name | No. samples | No. taxa | Host | Description | Reference |
| --- | --- | --- | --- | --- | --- |
| Human gut | 318 | 8,374 | Human | Stool microbiome of subject A at the OTU level | [3] |
| Human palm | 133 | 1,294 | Human | Left palm microbiome of subject F4 at the genus level | [4] |
| Human tongue | 135 | 372 | Human | Tongue microbiome of subject F4 at the genus level | [4] |
| Mouse gut | 30 | 6,285 | Mouse | Stool microbiome of Mice at the OTU level | [5] |
| Marine plankton bacteria | 90 | 28,951 | N/A | Bacteria relative abundance of Martin Platero at the OTU level | [6] |
| Marine plankton eukarya | 93 | 21,044 | N/A | Eukarya relative abundance of Martin Platero at the OTU level | [6] |

**TABLE S2.**
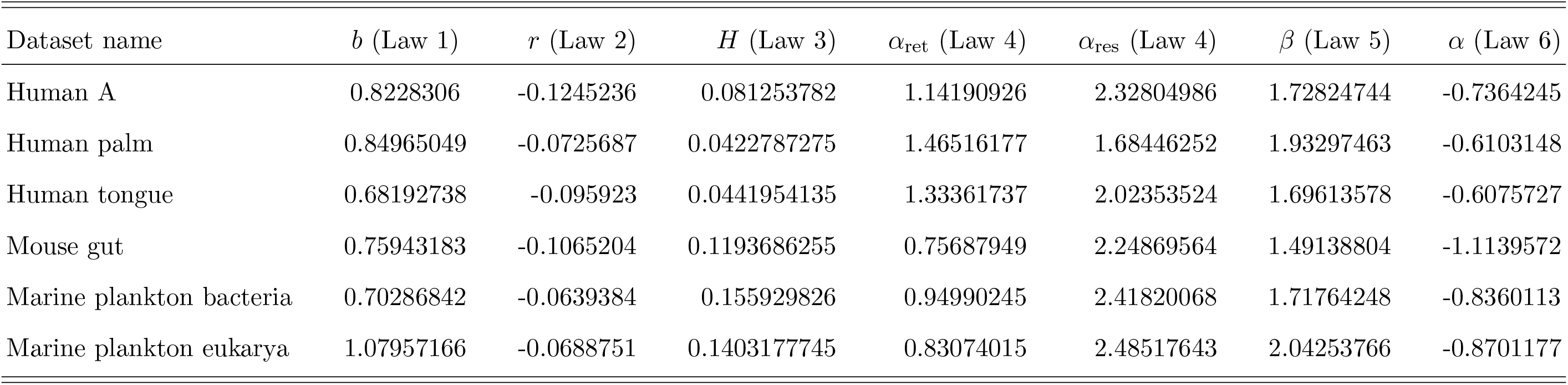
The exponents of scaling laws observed in the original time series.

| Dataset name | $b$ (Law 1) | $r$ (Law 2) | $H$ (Law 3) | $\alpha_{\text{ret}}$ (Law 4) | $\alpha_{\text{res}}$ (Law 4) | $\beta$ (Law 5) | $\alpha$ (Law 6) |
| --- | --- | --- | --- | --- | --- | --- | --- |
| Human A | 0.8228306 | -0.1245236 | 0.081253782 | 1.14190926 | 2.32804986 | 1.72824744 | -0.7364245 |
| Human palm | 0.84965049 | -0.0725687 | 0.0422787275 | 1.46516177 | 1.68446252 | 1.93297463 | -0.6103148 |
| Human tongue | 0.68192738 | -0.095923 | 0.0441954135 | 1.33361737 | 2.02353524 | 1.69613578 | -0.6075727 |
| Mouse gut | 0.75943183 | -0.1065204 | 0.1193686255 | 0.75687949 | 2.24869564 | 1.49138804 | -1.1139572 |
| Marine plankton bacteria | 0.70286842 | -0.0639384 | 0.155929826 | 0.94990245 | 2.41820068 | 1.71764248 | -0.8360113 |
| Marine plankton eukarya | 1.07957166 | -0.0688751 | 0.1403177745 | 0.83074015 | 2.48517643 | 2.04253766 | -0.8701177 |

**TABLE S3.** The exponents of scaling laws observed in the shuffled time series.

| Dataset name | $b$ (Law 1) | $r$ (Law 2) | $H$ (Law 3) | $\alpha_{\text{ret}}$ (Law 4) | $\alpha_{\text{res}}$ (Law 4) | $\beta$ (Law 5) | $\alpha$ (Law 6) |
| --- | --- | --- | --- | --- | --- | --- | --- |
| Human A | 1.16000329 | -0.1378984 | 0.001054554 | 1.10798649 | 2.44986712 | 1.72824744 | -0.7247134 |
| Human palm | 1.00759097 | -0.0576791 | -0.0068694355 | 1.47452508 | 1.81249994 | 1.93297463 | -0.595858 |
| Human tongue | 0.87138121 | -0.1171765 | 0.000199442 | 1.31220778 | 2.14643671 | 1.69613578 | -0.5938549 |
| Mouse gut | 0.85911145 | -0.1936557 | -0.021825658 | 0.77327667 | 2.18637321 | 1.49138804 | -1.098362 |
| Marine plankton bacteria | 1.3017074 | -0.0888345 | 0.0138457875 | 0.88909178 | 2.5702022 | 1.71764248 | -0.7707422 |
| Marine plankton eukarya | 1.84317825 | -0.0811869 | 0.0158372265 | 0.79562728 | 2.69942275 | 2.04253766 | -0.7934994 |

**FIG. S1.**
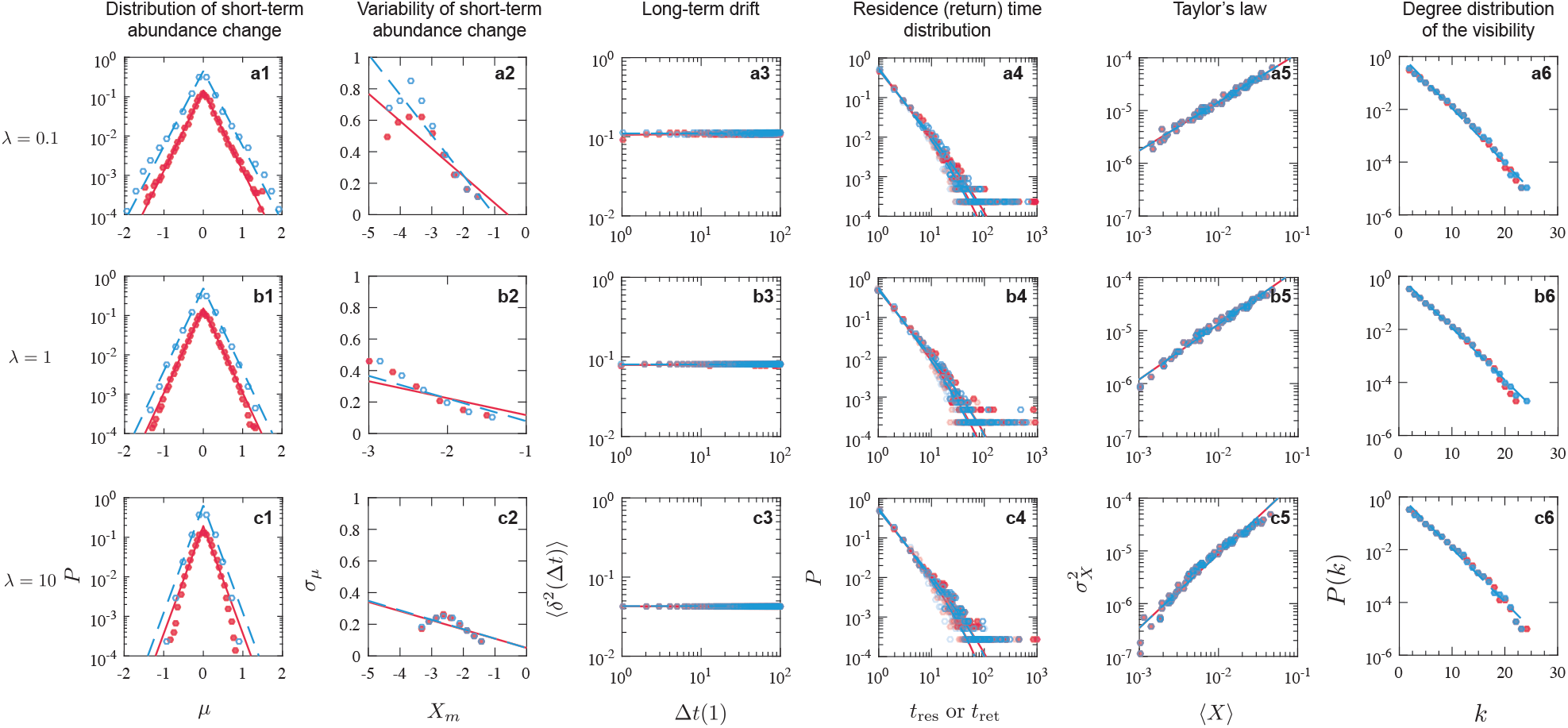
Scaling laws observed from simulated time series generated by the stochastic GLV model with different immigration rates. In the main manuscript, we chose the immigration rate λ = 0. Here, we showed the scaling laws with different immigration rates: λ = 0.1 (row 1); λ = 1 (row 2) and λ = 10 (row 3). *η* = 1.205, *m* = 7.9158, *C* = 1, *σ* = 0.1 and *N* = 100.

**FIG. S2.**
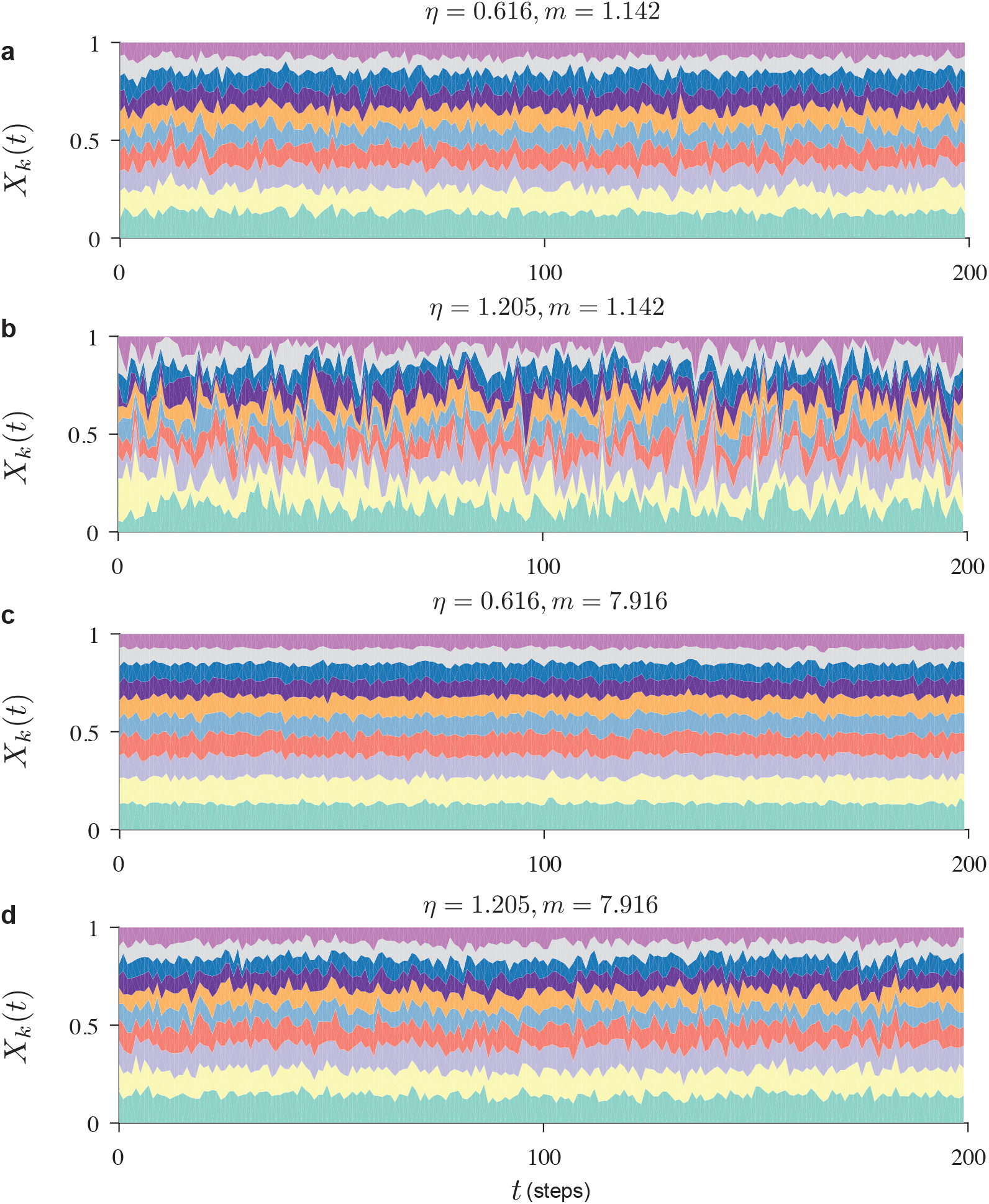
Time series generated from the stochastic GLV model with various parameter combinations. (a) *η* = 0.616, *m* = 1.142; (b) *η* = 1.205, *m* = 1.142; (c) *η* = 0.616, *m* = 7.916 and (d) *η* = 1.205, *m* = 7.916. *C* = 1, *σ* = 0.1, and N = 100. Stream widths reflect the relative abundances of those species. The total simulation step is 1000, d*t* = 0.01 and we showed the abundances of most abundant 10 species during the last 200 steps.

**FIG. S3.**
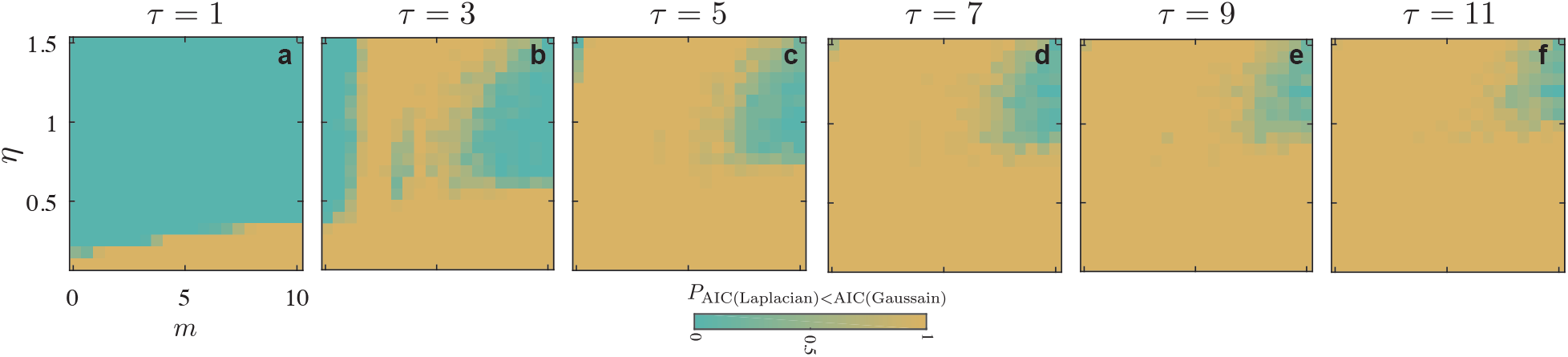
The distribution of short-term abundance change depends on the time span. The population growth rate is defined as: *μ*_*k*_(t) ≡ log(*X*_*k*_(*t* + *τ*)/*X*_*k*_(*t*)), where *τ* is the time span of the “short term”. (a) *τ* = 1; (b) *τ* = 3; (c) *τ* = 5; (d) *τ* = 7; (e) *τ* = 9 and (f) *τ* = 11. C = 0.4, *σ* = 0.05. The color in the phase diagram has the same meaning as shown in Fig.3 in main text. The time step is d*t* = 0.01, total time *T* = 1000, and the number of species is *N* = 100.

**FIG. S4.**
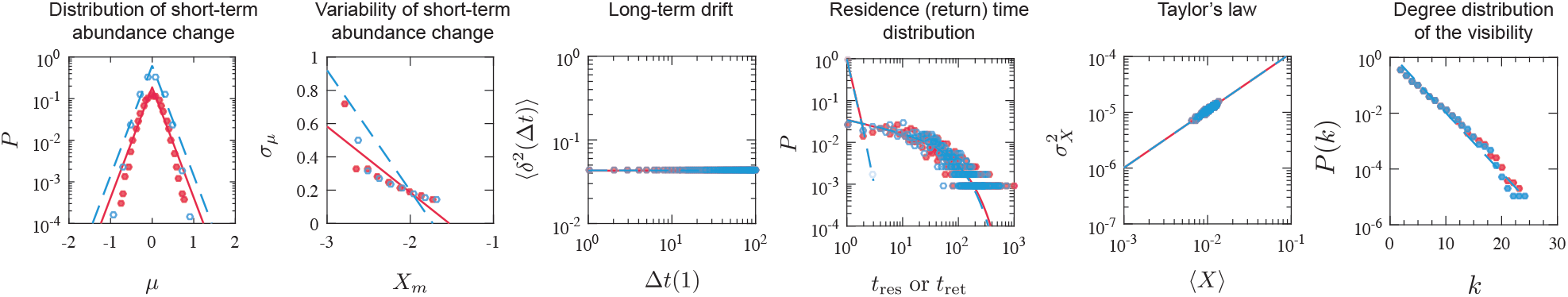
Scaling laws observed from simulated time series generated by the stochastic GLV model without inter-species interactions. *η* = 1.205, *m* = 7.9158, *C* = 0, *σ* = 0, and *N* = 100.

**FIG. S5.**
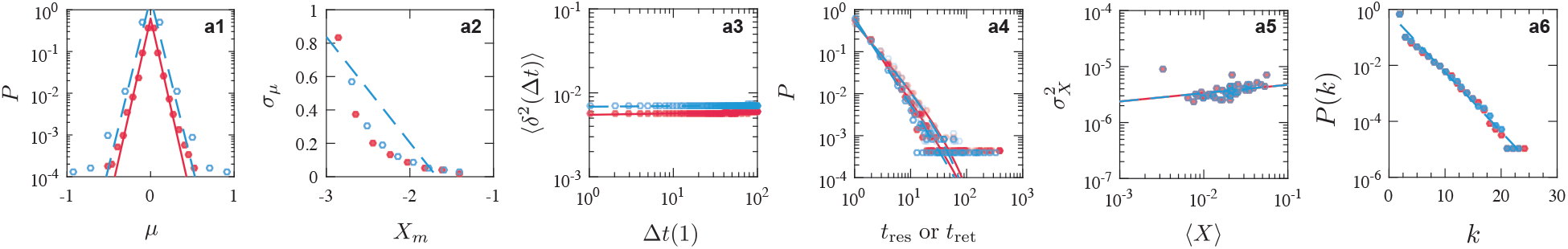
Scaling laws observed from simulated time series generated by the stochastic GLV model with nonlinear multiplicative noise. *η* = 1.205, *m* = 7.9158, *C* = 1, *σ* = 0.1, and *N* = 100.

